# Identification of potential inhibitors against Inosine 5’-Monophosphate Dehydrogenase of *Cryptosporidium parvum* through an integrated in silico approach

**DOI:** 10.1101/2024.09.22.614371

**Authors:** Abdullah Al-Mamun, Shah Imran Hossain, Abu Tayab Moin, Md Shafiqul Islam Rakib, Md Mahedi Hasan, Eshma Binte Yousuf, Shams Nur Powshi, Enayetul Islam, Nasrin Jahan Salma Tumpa, Asmaul Hosna, Dil Umme Salma Chowdhury, Mohabbat Hossain, Sabrina Shameen Alam, Nazneen Naher Islam

## Abstract

The protozoan parasite *Cryptosporidium*, found in several vertebrates, including humans, is the source of the global infection known as cryptosporidiosis, which manifests as acute gastroenteritis, abdominal pain, and diarrhea. Although infections in certain individuals have been linked to other species, *Cryptosporidium parvum* is the main cause of illnesses in humans. Lactate Dehydrogenase, Inosine 5′-Monophosphate Dehydrogenase (IMPDH), and several other targets have been identified by the genome sequencing of *C. parvum*. Bioactive phytochemicals derived from nature have enormous potential as anti-cryptosporidiosis agents. The study aimed to identify new anti-cryptosporidial agents that work against the IMPDH of the parasite by using integrated in silico approaches. In this study, a total of 24 bioactive phytochemicals were screened virtually through molecular docking and ADMET (Absorption, Distribution, Metabolism, Excretion, and Toxicity) analyses. Four lead compounds were identified, including Brevelin A (−8.9 kcal/mol), Vernodalin (−8.7 kcal/mol), Luteolin (−8.6 kcal/mol), and Pectolinarigenin (−8.1 kcal/mol), against the IMPDH protein (PDB ID: 4IXH) from the parasite. All the lead compounds had excellent pharmacokinetic and pharmacodynamic characteristics. The toxicity analysis showed satisfactory results with no major side effects. All of the selected compounds showed no violation of Lipinski’s rules of five, indicating the possibility of oral bioavailability as potential drug candidates. In the majority of cases, target class prediction-identified enzymes, as well as investigational and experimental drugs, have been found to have structural similarities to the lead compounds. With significant biochemical interactions, all of the targeted phytochemical compounds have demonstrated excellent pharmacokinetics and better bioavailabilities. The findings strongly recommend in vitro experimental studies to aid in the development of novel therapeutics against *Cryptosporidium parvum*.

## 1. Introduction

Diarrhea is the most prevalent cause of mortality among young infants worldwide. Many species of *Cryptosporidium* account for a major part of the diarrheal burden globally [1]. Enteric protozoan parasites, or *Cryptosporidium spp*., are widespread around the world. They are abundant in the environment and can infect a variety of species, including humans that are immunocompetent or immunocompromised. It is becoming more widely acknowledged that they can trigger both sporadic infections and more extensive food and waterborne outbreaks [2, 3].

Although there are 26 known species of intestinal protozoa called *Cryptosporidium spp.,* people are most frequently infected by *C. hominis* and *C. parvum*. The hallmark of an infection is copious, watery diarrhea. In immune-competent adults, the illness is self-limiting, but in immunocompromised individuals and small children, it may be linked to fulminant sickness. Infection with *Cryptosporidium* has been linked to prolonged diarrhea and 2-3 times greater mortality in young children [4–7]. Along with outbreaks and cases of diarrhea in developed countries, *Cryptosporidium* is the cause of diarrhea and malnutrition in young infants in developing countries. The majority of deaths from diarrheal disease occur in Sub-Saharan Africa and South Asia, due to the prevalence of *Cryptosporidium* diarrheal disease. *Cryptosporidium* is the most common cause of diarrhea in children under 24 months old. Although the clinical manifestations of diarrhea caused by *Cryptosporidium* peaked between the ages of six and eleven months, infections started in the in the initial stages of life. The number of diarrhea episodes in children aged 24 months caused by *Cryptosporidium* is estimated to be 2.9 million in Sub-Saharan Africa and 4.7 million in the South Asian region, which includes Bangladesh, India, Pakistan, Afghanistan, and Nepal. In both regions, *Cryptosporidium* is responsible for about 202,000 deaths annually; deaths from cases that are attributable to the parasite are 59,000 higher than deaths from cases that are not [8].

*Cryptosporidium*, an enteropathogen, is highly prevalent in underdeveloped nations, affecting young children. Patients with compromised immune systems, such as those with HIV/AIDS, face significant challenges due to malnourishment, chronic physical inactivity, and other health-related issues [9]. The fecal-oral route is the main way that *Cryptosporidium* spreads. It can be acquired indirectly through contaminated food or water or directly from an infected person or animal. Feces from cattle and other animals can contaminate crops, agricultural goods, and surface water, causing zoonotic transmission. Several of the parasite’s intrinsic characteristics account for its epidemiologic behavior. Oocysts can spread for up to two months after diarrhea has stopped. They are contagious as soon as they are expelled in feces and shed in large quantities (up to 109 per stool) [10].

Certain strains of *C. parvum* and *C. hominis* have low infectious doses; symptoms have been reported in healthy volunteers after as little as 9–10 oocysts. Oocysts that are kept moist can withstand chlorination and remain infectious for a minimum of six months in the environment. They can also withstand more than ten days in fully chlorinated recreational water venues. Transmission can go on for days before public health officials become aware of an outbreak because of the lengthy incubation period (average 7 days, range 1–30 days). The most vulnerable are those who lack protective immunity, such as children living in endemic areas and immunocompromised individuals, as evidenced by age-related declines in illness incidence. Immunity gained through prior exposure is therefore protective [11–13].

Cryptosporidiosis is caused by the coccidian *Cryptosporidium*, a parasite that infects epithelial cells of several vertebrate hosts. Mammals, including humans and domestic animals, are susceptible to *C. parvum* infections of the gastrointestinal tract, which can result in acute watery diarrhea and weight loss. Hosts lacking cells of immune system such as CD4+ T are more sensitive to parasite infection and have a greater possibility to have severe, potentially deadly consequences. The control of the immune system is essential in the battle against infection. A study employing mouse infection models suggests that IFN-g is important for both the eradication of infection via CD4+ T-cells and the largely preventive innate immune response against infection were observed in immunocompromised mice. At least in part, CD4+ lymphocytes in the intraepithelial layer play an important role in regulating cryptosporidial infection at the mucosal level by generating IFN-g., which directly prevents the formation of parasites in enterocytes. An inflammatory response is brought on by ruminant infection, resulting in an increase in the quantity of various T-cell subpopulations showing up in the villi. Furthermore, infection causes the intestine to express more pro-inflammatory cytokines such as TNF-a, IFN-g, and IL-12. Since these type of cytokines tend to be crucial in the pathogenesis of inflammatory bowel disease, it is possible that they are involved in the mucosal etiology of cryptosporidiosis [14]. In the first year of life, *Cryptosporidium* infections affected about 40% of children in Bangladesh [15]. It has been determined that breastfeeding and the IgA in breast milk are protective factors. *Cryptosporidium* infection has been associated with HLA class II alleles and mutations in the mannose-binding lectin gene, thus host genetic vulnerability is also implicated [16–18].

In the past, a number of studies on the treatment of *Cryptosporidium* have been conducted. Numerous studies have demonstrated that nitazoxanide enhances clinical response and shortens the duration of diarrhea and oocyst shedding in immunocompetent adults and children with cryptosporidiosis [19, 20]. Nitazoxanide is the sole medicine approved by the US Food and Drug Administration for the therapeutic management of cryptosporidiosis in people who are immunocompetent, however it only works occasionally. Nevertheless, there is no proven cure for this fatal illness that affects newborns, immunocompromised individuals, and neonatal cattle. Because *C. parvum* lacks both oxidative phosphorylation and the Krebs cycle, it can only produce metabolic energy through glycolysis [21].

In this instance, naturally occurring bioactive compounds can be a highly important and promising source for developing novel anti-cryosporidiosis medications. Alkaloids, terpenoids, coumarins, flavonoids, nitrogen-containing compounds, organosulfur compounds, phenolics, etc. are examples of bioactive compounds found in plants. These substances show a broad range of bioactivities, including anti-inflammatory, immunostimulatory, anticancer, antioxidant, and antimicrobial properties. Compared to the conventional approach, structure-based drug design is quickly becoming a vital tool for quicker and more affordable lead discovery. So, the study aimed to identify possible inhibitors as a novel drug for diarrheal diseases caused by *Cryptosporidium parvum*, targeting its Inosine 5′-Monophosphate Dehydrogenase enzyme by applying integrated *in silico* approaches.

## 2. Methods

**Fig 1** demonstrates the layout of the study.

**Fig 1.**
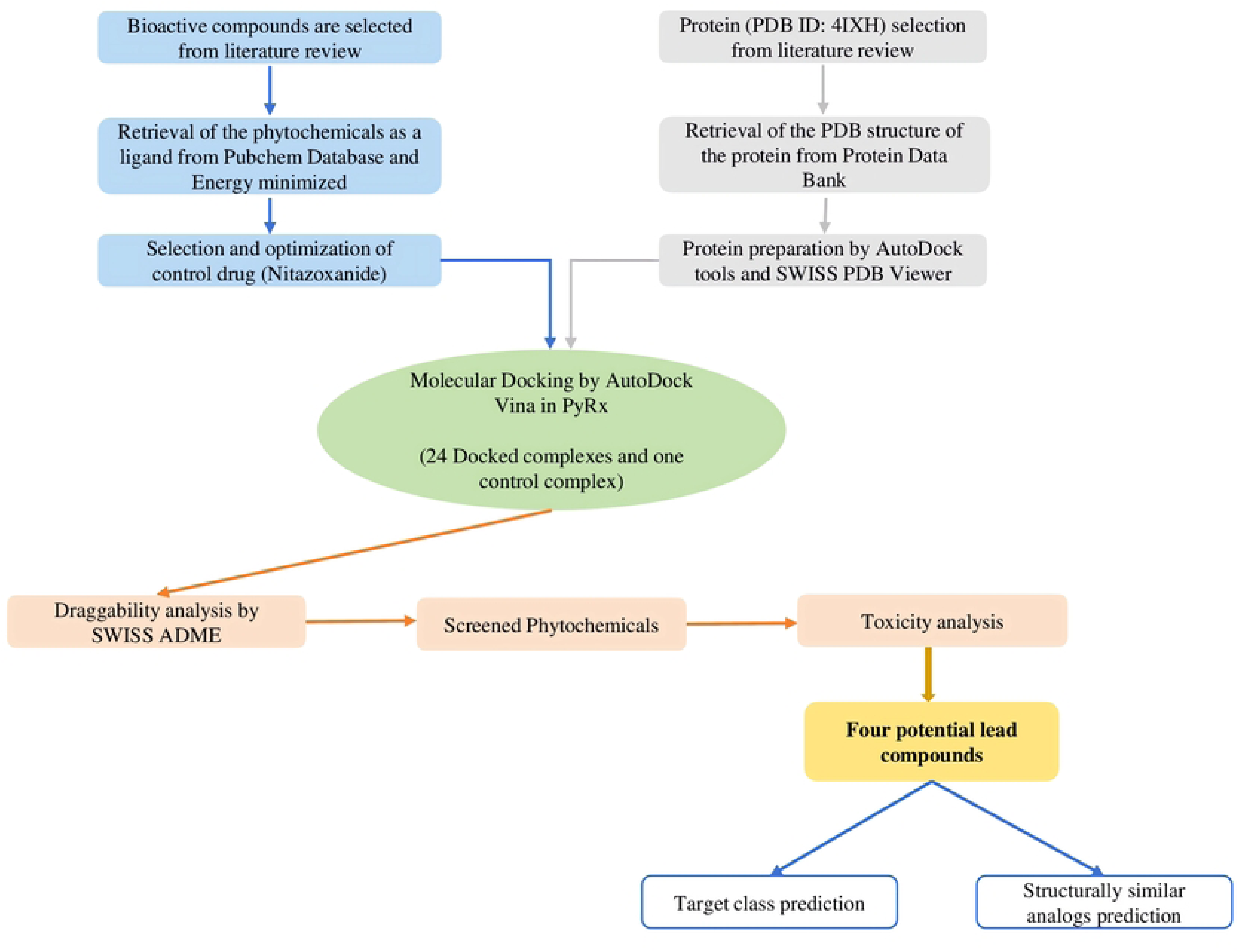
Simplified overview of the multiple steps carried out in the present experiment for a virtual evaluation of biologically active phytochemicals against cryptosporidiosis.

### 2.1. Receptor preparation

The three-dimensional (3D) structure of the targeted pathogenic protein of *Cryptosporidium parvum* was retrieved by searching the RCSB protein databank (https://www.rcsb.org) with the protein data bank ID (PDB ID) ‘4IXH’. The protein 4IXH contained 1,444 amino acids, with a resolution of 2.10 Å and a total structure weight of 157.37 kDa [22].

Using Biovia Discovery Studio Visualizer, the obtained PDB file for the protein was opened to exclude water molecules, metal ions, and co-factors. The AutoDock tool (version 1.5.7) was employed to add polar hydrogen, Kollman charges, and recover missing atomic residues [23]. The SWISS PDB Viewer (version 4.10) was used to utilizes the GROMOS96 43b1 force field, which is necessary to retain the protein structure, to ensure energy minimization [24, 25]. Protein-energy minimization has been effectively achieved using the GROMOS96 43b1 force field in previous studies [25–27].

### 2.2. Ligand preparation

To develop a list of phytochemicals and the control ligand, nitazoxanide, appropriate research papers were searched through PubMed, Google Scholar, and Scopus databases. Every phytochemical’s 3D conformers were obtained from the PubChem database (https://pubchem.ncbi.nlm.nih.gov/) in SDF (Structured Data Files) format. The Avogadro software (version 1.90.0) was then used to convert these conformers into PDB format [64]. Merck Molecular Force Field (MMFF94) optimized phytoligands [25, 28].

### 2.3. Molecular docking

Molecular docking is a valuable tool for determining interactions between ligands and targets by screening thousands of different molecules computationally. The ligands were docked to the target receptors and their binding affinities were assessed using the PyRx-free package 0.8 [25, 29]. In this strategy, ligands were maintained flexible, and proteins stiff. To complete intermediary tasks, such as generating grid boxes and PDBQT files for proteins and ligands, PyRx software in combination with AutoDock Vina was utilized [30, 64]. To find an unknown ligand-binding site in a target protein, this study used the blind docking strategy, which makes use of the entire protein surface. The docked complexes were then visualized using Discovery Studio for analyzing ligand-protein interactions [25, 64].

### 2.4. ADME analysis and toxicity prediction

Drug levels and the kinetics of drug interaction with tissues within an organism are influenced by four major factors: absorption, distribution, metabolism, and excretion (ADME). The pharmacological activity and efficacy of a drug are greatly influenced by these features [31, 64]. The SwissADME online tool (http://www.swissadme.ch/) was employed for predicting a variety of pharmacokinetic and pharmacodynamic parameters, including physiochemical features, lipophilicity, water solubility, drug-likeness, and medicinal chemistry properties. The blood-brain barrier (BBB) of the substances under research was examined utilizing the BOILED-Egg model [25, 32, 33].

The toxicity analysis was performed using the pkCSM server (https://biosig.lab.uq.edu.au/pkcsm/prediction), which uses a unique method of distance-based graph signatures to predict and improve the pharmacokinetic and toxicological features of small molecules [25, 64]. This server assessed the toxicity profiles of the top candidates using a variety of hazardous metrics, including hepatotoxicity, skin sensitization, hERG I and II inhibitory characteristics, LOAEL, LD50, and others [34].

### 2.5. Drug target prediction and ligand homology screening

Therapeutic target prediction confirms metabolic bioactivity. Choosing small molecules as targets makes it easier to assess drug safety and uncover new applications for treatment. The web-based tool SwissTargetPrediction (http://www.swisstargetprediction.ch/) was used in this work to identify the future macromolecular targets of drug candidates [25, 35]. The software predicts a library of 3,76,342 identified biologically active compounds on about 3068 proteins using a mix of 2D and 3D analogies. The SwissSimilarity server (http://www.swisssimilarity.ch/) was used to predict structural analogs from the DrugBank database that were comparable to the top candidates [25, 64]. These analogs could be repurposed for the treatment of diarrheal diseases. The server employs several techniques, including FP2 fingerprints, electroshape, spectrophores, shape-IT, and align-IT approaches, to carry out its prediction [36].

## 3. Results

### 3.1. Molecular docking assessments

The target protein was successfully docked with 24 phytochemicals (**Fig 2**), with binding affinities ranging from −5.5 kcal/mol to −10.2 kcal/mol. The control drug, Nitazoxanide, scored −8.1 kcal/mol. In comparison to the control, 8 phytochemicals (33.33%) with a score equal to or higher than the control were accepted, while 16 phytochemicals (66.67%) with a score lower than the control were rejected. Thiarubrine A (−5.5 kcal/mol) had the lowest binding affinity, and Epigallocatechin Gallate (−10.2 kcal/mol) had the highest binding affinity according to molecular docking analysis (**Table 1**). Eight of the highest-scoring phytochemicals underwent the ADMET assay, and only four of the best candidate compounds were chosen for additional analysis and molecular interactions. Brevelin A (−8.9 kcal/mol), Vernodalin (−8.7 kcal/mol), Luteolin (−8.6 kcal/mol), and Pectolinarigenin (−8.1 kcal/mol) (**Fig 3**) were the four leading candidates.

**Fig 2.**
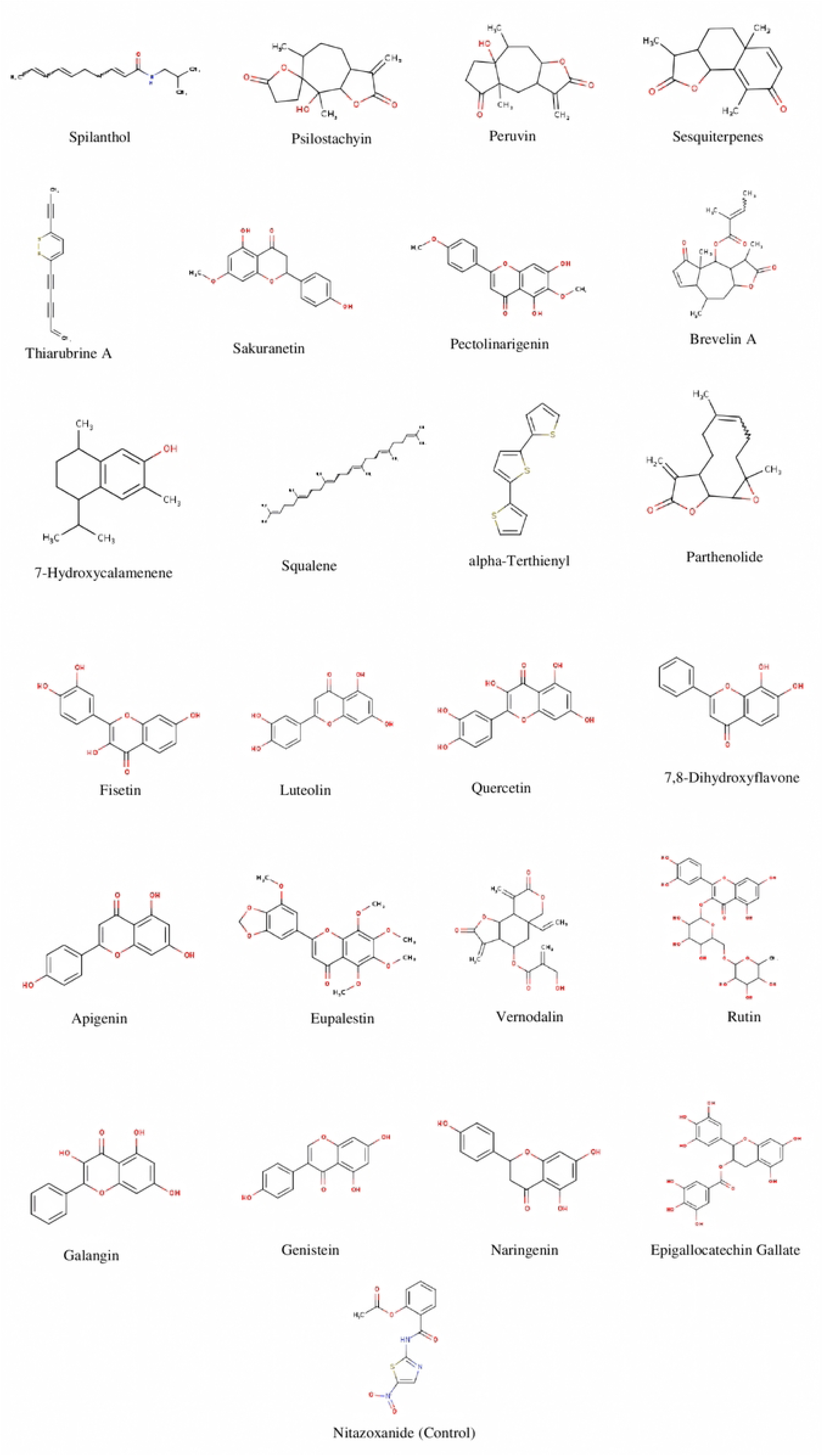
Two-dimensional chemical structures of 24 phytochemicals were employed for virtual evaluation, as well as the control compound Nitazoxanide.

**Fig 3.**
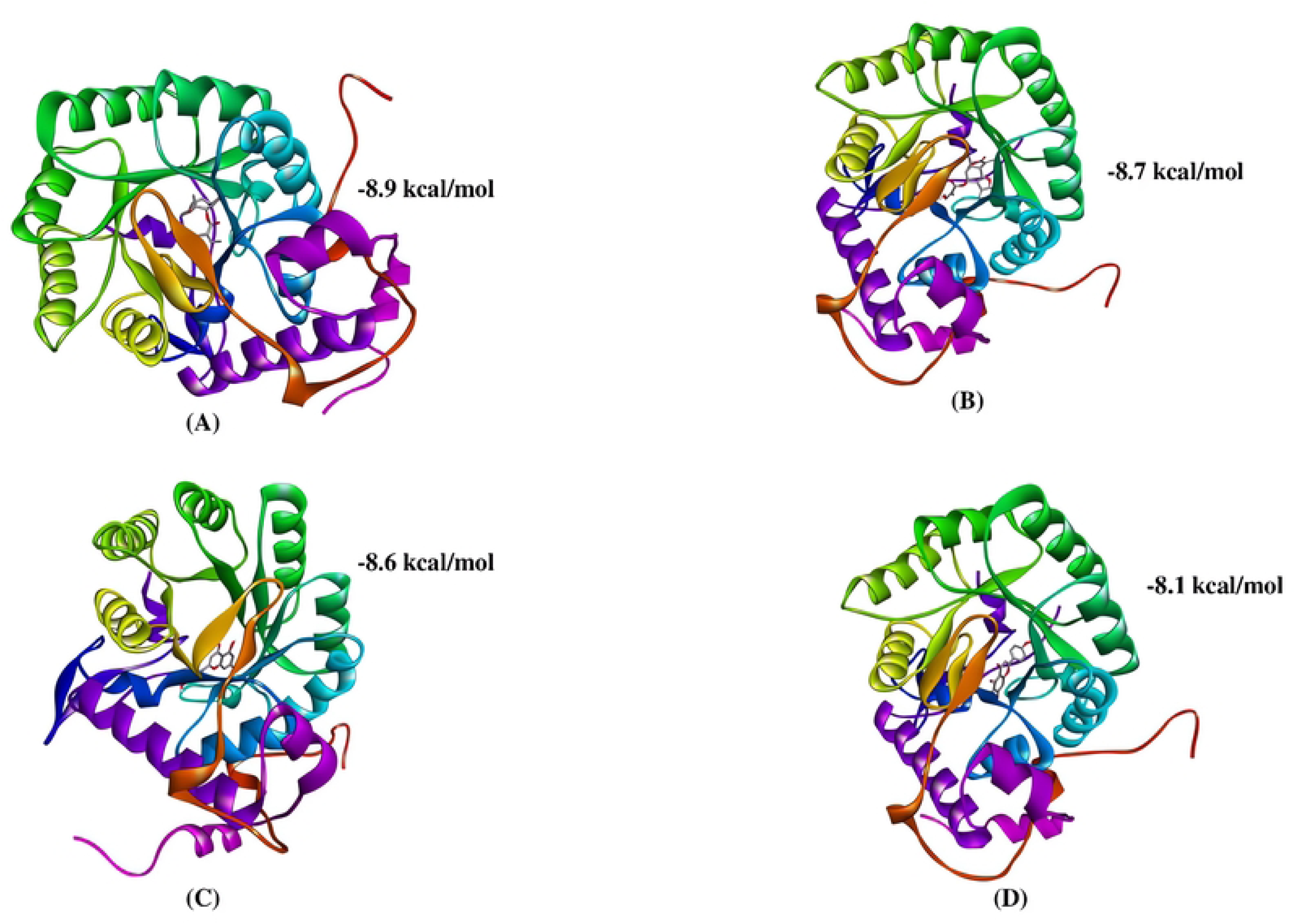
Ligand-protein interactions for the top four lead compounds, collectively with their docking scores, are depicted in ribbon view: (A) Brevelin A, (B) Vernodalin, (C) Luteolin, and (D) Pectolinarigenin.

**Table 1.**
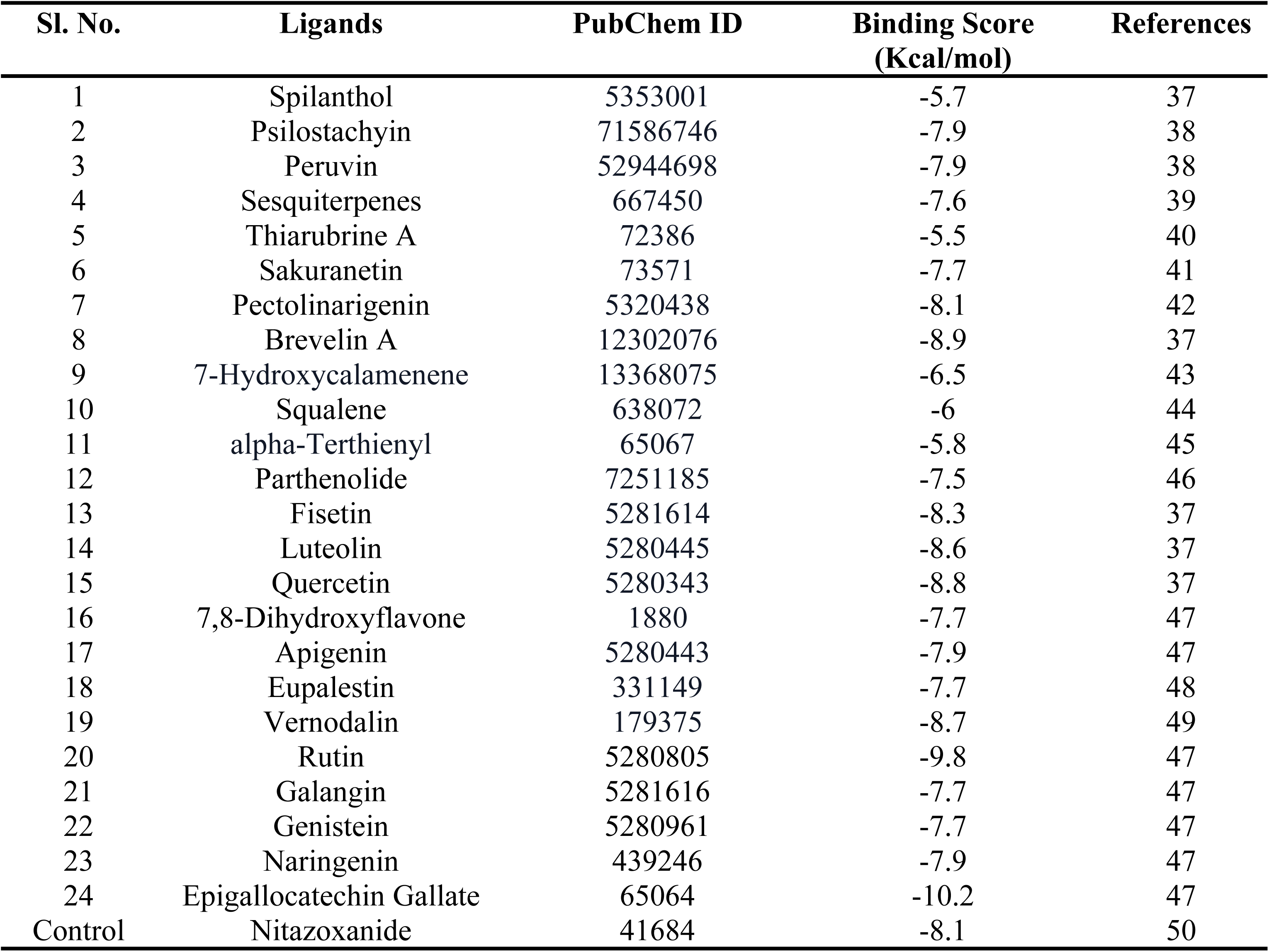
Phytochemicals with docking scores (−Kcal/mol) as well as the control compound Nitazoxanide.

### 3.2. Molecular interactions of the lead compounds

Among the four lead compounds, three (Vernodalin, Luteolin, and Pectolinarigenin) exhibited both hydrogen bonds and hydrophobic interactions (**Table 2**). There were no hydrophobic interactions in Brevelin A. It showed interactions with Thr221, Gly214, and Lys210. Pectolinarigenin contained the most usual hydrogen bonds. It interacted with Ser217, Gly275, Ser48, Met273, Asp252, Gly303, Met50, Cys219, and Ile255. Of these residues, Met50, Cys219, and Ile255 were found in hydrophobic interactions, whereas the remaining residues were involved in typical hydrogen bonding. In the case of luteolin, Cys219 is involved in hydrophobic interaction, on the other hand, Gly275 and Ser164 showed hydrogen bonding. Vernodalin interacted with Gly214, Gly303, Ser48, and Met273 with conventional hydrogen bonds. It showed interactions with Ile213, Met302, and Met50 with hydrophobic interactions (**Figs 4 and 5**).

**Fig 4.**
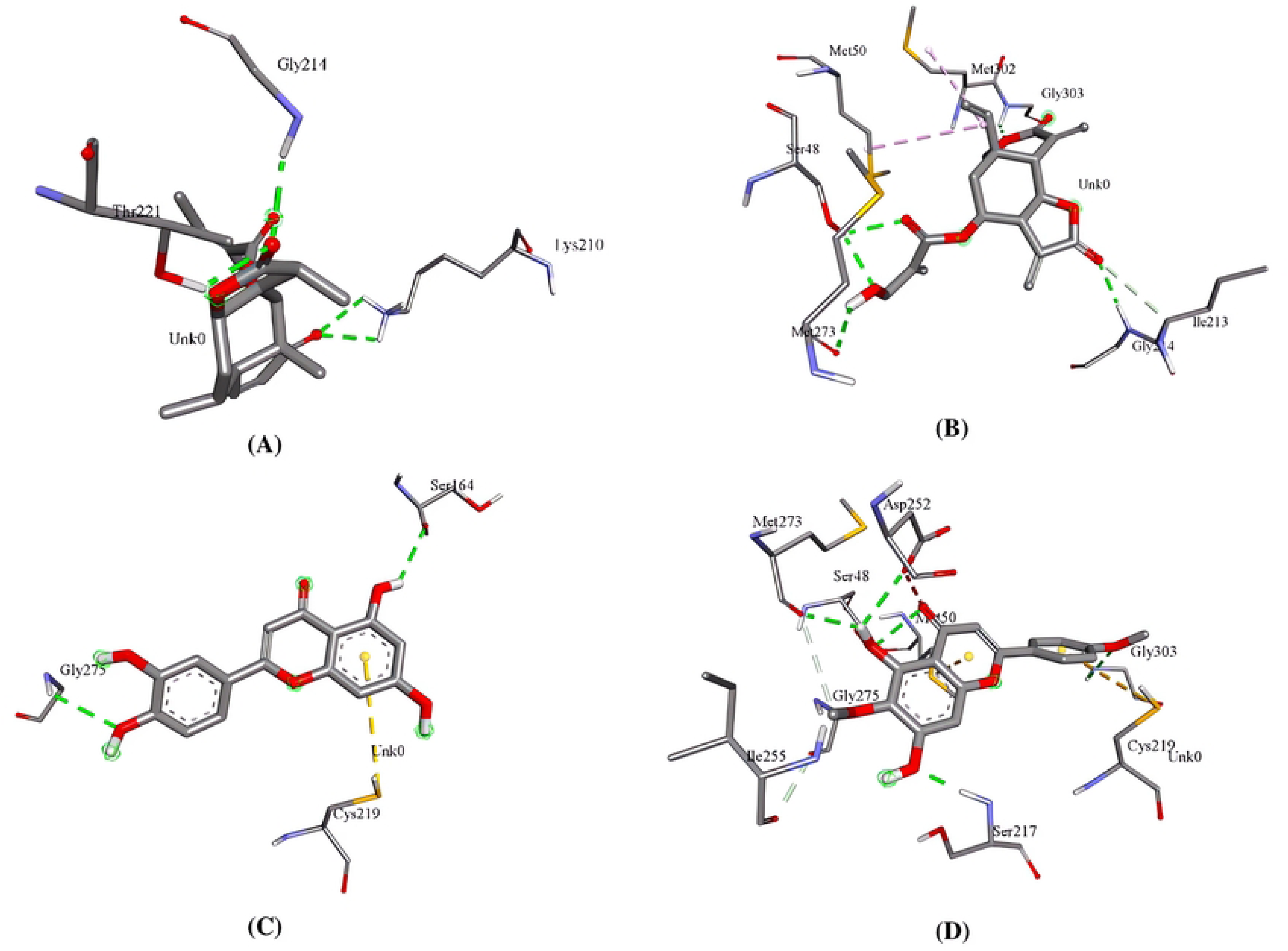
Three-dimensional molecular interaction systems displaying numerous interacting residues of the protein macromolecule with (A) Brevelin A, (B) Vernodalin, (C) Luteolin, and (D) Pectolinarigenin.

**Fig 5.**
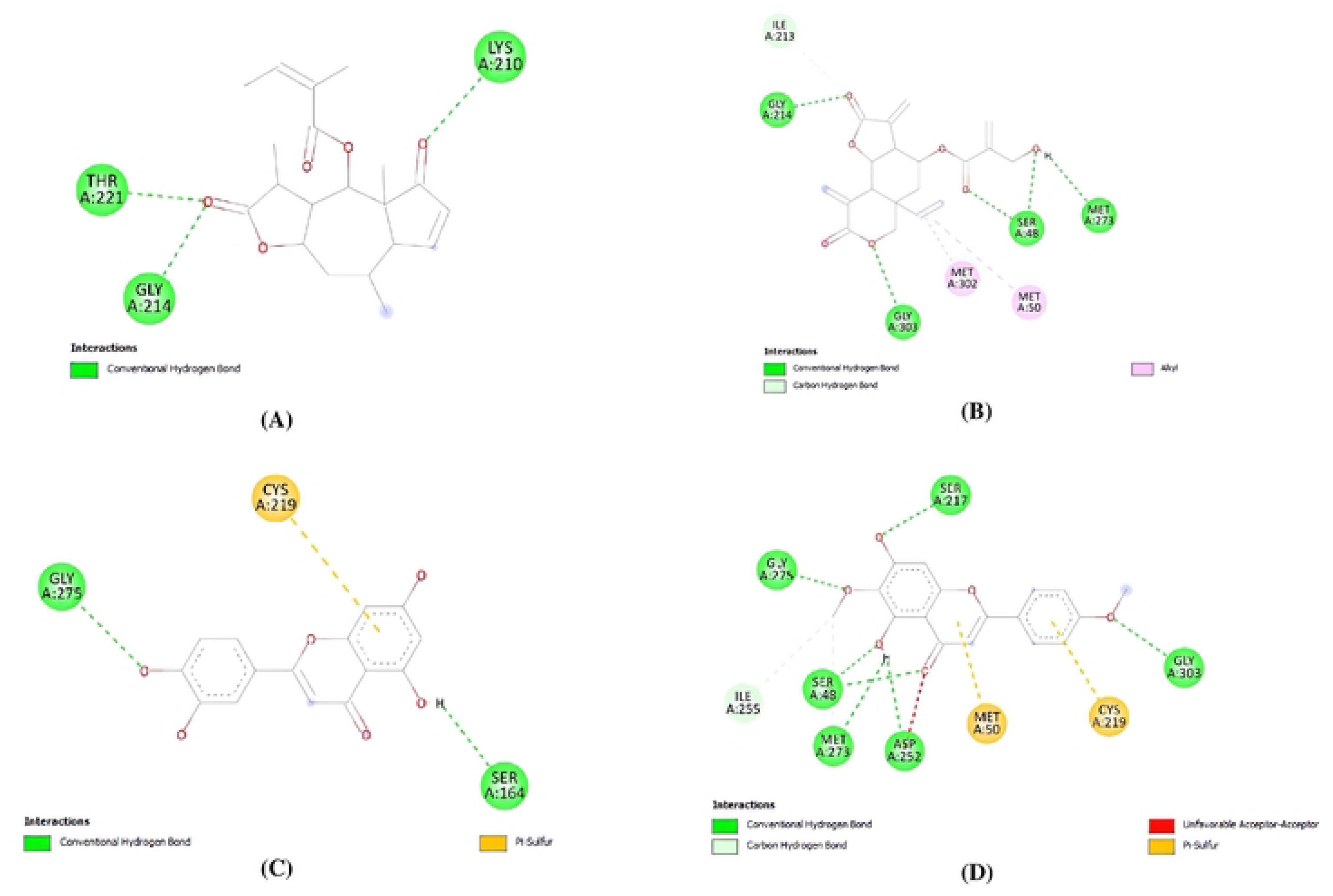
Two-dimensional molecular interaction systems displaying numerous interacting residues of the protein macromolecule with (A) Brevelin A, (B) Vernodalin, (C) Luteolin, and (D) Pectolinarigenin.

**Table 2.** Analysis of the top four phytochemicals’ interaction sites and binding affinities.

| Macromolecules | Ligands | Ligands binding sites | Residues involved in hydrogen bond formation (Distance in Å) | Number of hydrogen bonds formed | Residues involved in hydrophobic interactions | Binding affinity (kcal/mol ) |
| --- | --- | --- | --- | --- | --- | --- |
| 4IXH | Nitazoxanide (Control) | Gly303<br>Met302<br>Ser217<br>Gly254<br>Ser276<br>Asp163<br>Met50 | Gly303<sup>(2.64)</sup><br>Met302<sup>(2.39)</sup><br>Ser217<sup>(2.06)</sup><br>Gly254<sup>(2.01)</sup><br>Ser276<sup>(2.16)</sup> | 5 | Asp163<br>Met50 | -8.1 |
|  | Brevelin A | Thr221<br>Gly214<br>Lys210 | Thr221<sup>(2.25)</sup><br>Gly214<sup>(2.30)</sup><br>Lys210<sup>(2.69)</sup><br>Lys210<sup>(2.76)</sup> | 4 | - | -8.9 |
|  | Vernodalin | Gly214<br>Gly303<br>Ser48<br>Met273<br>Ile213<br>Met302<br>Met50 | Gly214(2.25 )<br>Gly303(2.85 )<br>Ser48(2.29)<br>Ser48(2.30)<br>Met273(2.52 ) | 5 | Ile213<br>Met302<br>Met50 | -8.7 |
|  | Luteolin | Gly275<br>Ser164<br>Cys219 | Gly275<sup>(2.70)</sup><br>Ser164<sup>(2.46)</sup> | 2 | Cys219 | -8.6 |
|  | Pectolinarigenin | Ser217<br>Gly275<br>Ser48<br>Met273<br>Asp252<br>Gly303<br>Met50<br>Cys219<br>Ile255 | Ser217<sup>(1.94)</sup><br>Gly275<sup>(1.81)</sup><br>Ser48<sup>(2.03)</sup><br>Ser48<sup>(2.69)</sup><br>Met273<sup>(2.32)</sup><br>Asp252<sup>(2.53)</sup><br>Gly303<sup>(2.81)</sup> | 7 | Met50<br>Cys219<br>Ile255 | -8.1 |

### 3.3. ADME analysis of top drug candidates

To evaluate the drug properties and drug-likeness of the leading drug candidates, different ADME features were calculated, including physicochemical parameters, pharmacokinetics, lipophilicity, water solubility, and medicinal chemistry. The results of the analysis of the inhibitory outcome with various CYP isoforms (CYP1A2, CYP2D6, CYP2C9, CYP2C19, CYP3A4) showed that while Luteolin and Pectolinarigenin exhibited interaction potentials with certain cytochrome P450 isoforms, Brevelin A and Vernodalin did not interact with any of them. Each of the most suitable candidates had substantial GI absorption. Brevelin A was found to have positive blood brain barrier permeability, while vernodalin, luteolin, and pectolinarigenin had negative blood brain barrier permeability. Good solubility properties were demonstrated by all four of the selected candidates. These the compounds complied Lipinski’s rule of five with 0 infractions. Furthermore, for each tested top-selected drug, the Ghose, Veber, Egan, and Muegge parameters were determined to be appropriate (0 violations). In terms of medicinal chemistry properties, Vernodalin displayed one violation, but Brevelin A, Luteolin, and Pectolinarigenin showed promising lead likeness properties. Brevelin A performed the best in terms of synthetic accessibility properties out of the four candidates (**Table 3 and Fig 6**).

**Fig 6.**
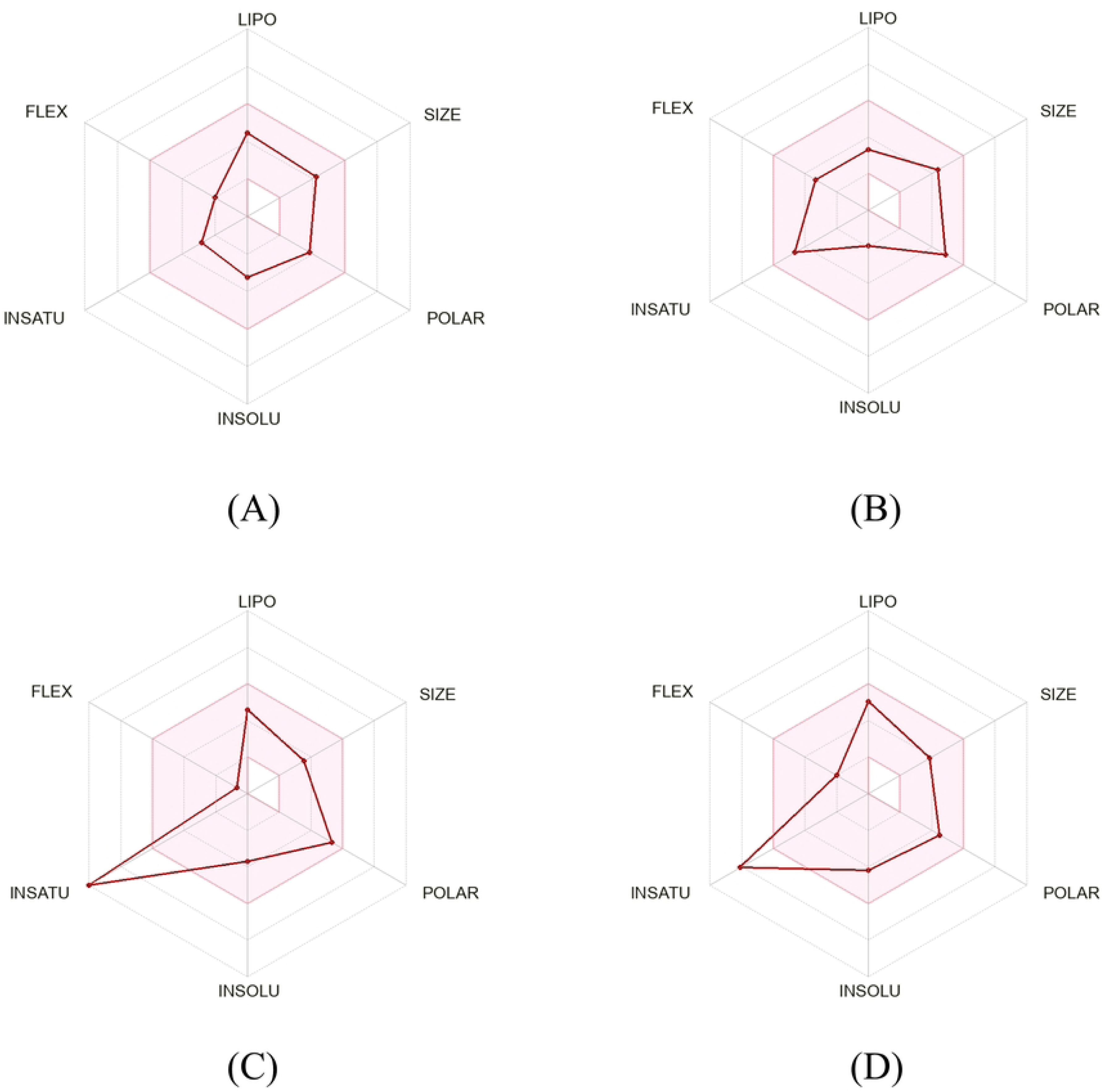
SwissADME radar displays multiple pharmacokinetic and pharmacodynamic parameters for (A) Brevelin A, (B) Vernodalin, (C) Luteolin, and (D) Pectolinarigenin. LIPO is for Lipophilicity, POLAR is for Polarity, INSOLU is for Insolubility, INSATU is for Insaturation, FLEX is for Flexibility.

**Table 3.**
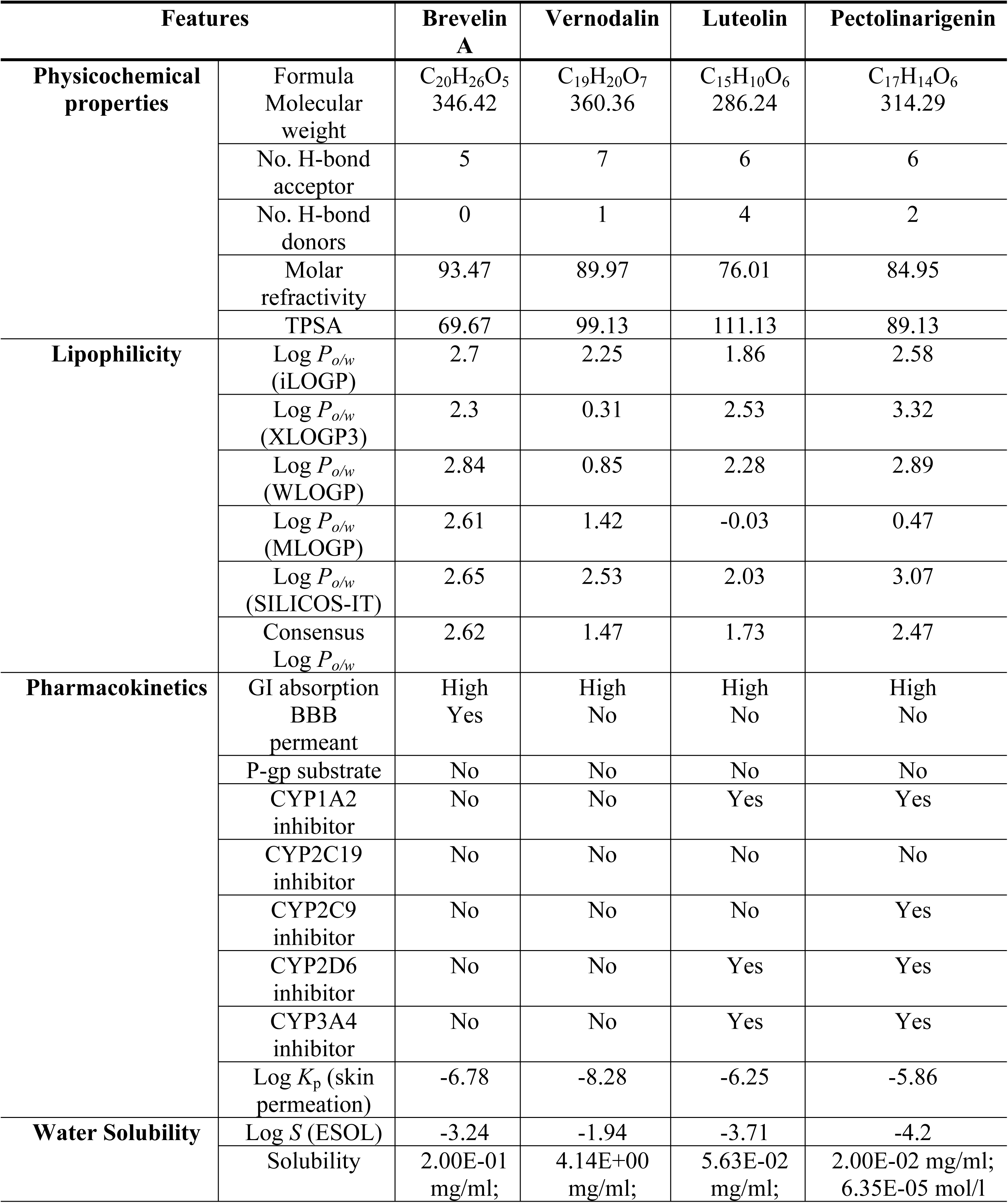
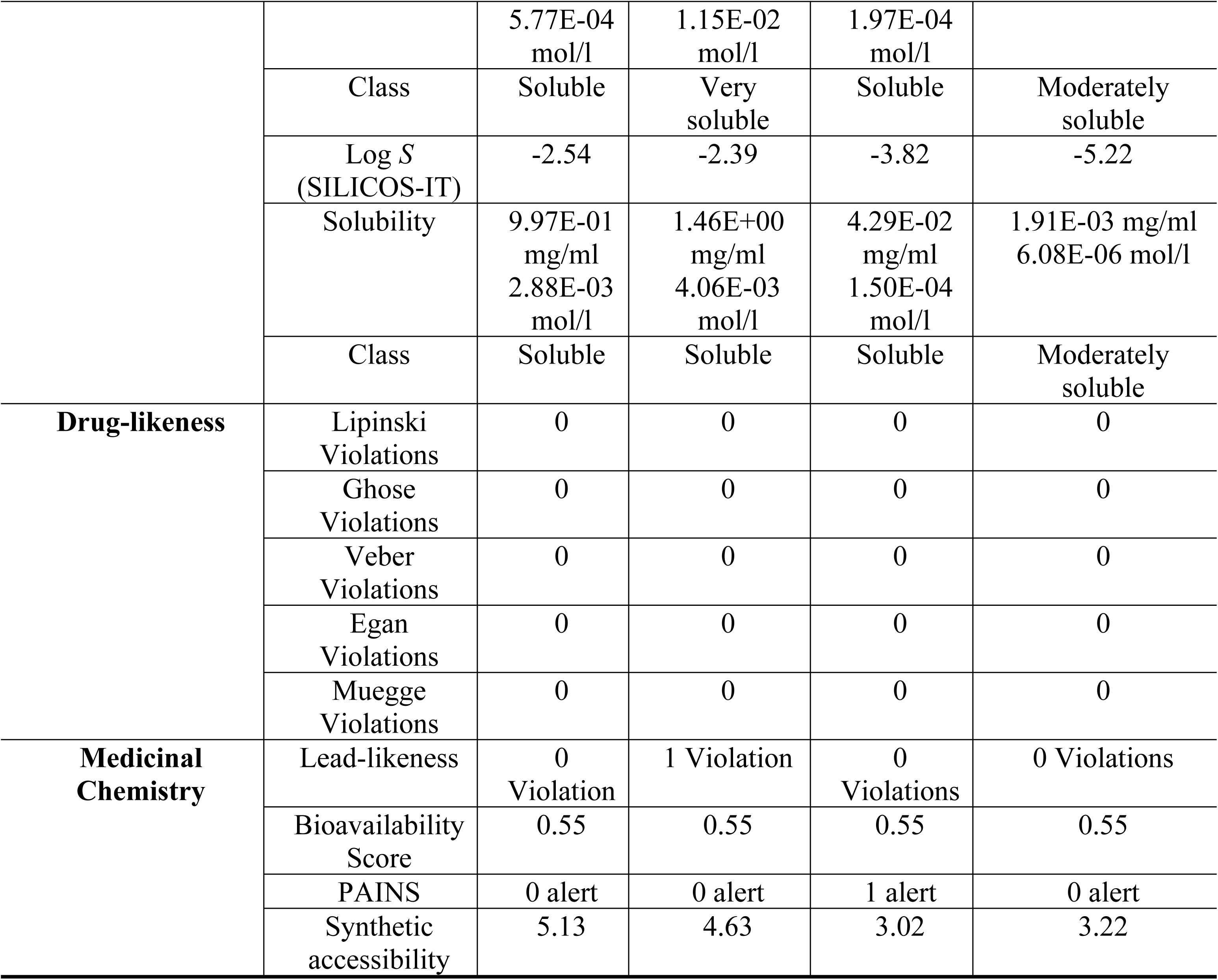
Evaluation of drug profile via ADME analyses for the top four lead compounds.

### 3.4. Toxicity profile examination of top drug candidates

A number of toxicity parameters were predicted using the pkCSM server, including hepatotoxicity, minnow toxicity, AMES toxicity, oral rat acute toxicity (LD50), oral rat chronic toxicity (LOAEL), and Tetrahymena pyriformis toxicity. Vernodalin exhibited AMES toxicity, while Brevelin A, Luteolin, and Pectolinarigenin did not exhibit any toxicity. None of the chosen candidates exhibited characteristics of hERG I and II inhibition. All the compounds exhibited negative results in the skin sensitization parameter and no effects in the hepatotoxicity criterion. Luteolin and Pectolinarigenin have a maximum tolerated dose of approximately 0.50 mg/kg/day for humans. The LD50 values of the four ligands didn’t show any notable toxicity. All of the candidates had minnow toxicity values greater than −0.3 log mM, indicating non-toxic potential (**Table 4**).

**Table 4.** Toxicity profile evaluation of assessed drug candidates.

| Model Name | Predicted Value |  |  |  |
| --- | --- | --- | --- | --- |
|  | Brevelin A | Vernodalin | Luteolin | Pectolinarigenin |
| AMES toxicity | No | Yes | No | No |
| Max. tolerated dose (human) | -0.081 mg/kg/day | 0.236 mg/kg/day | 0.499 mg/kg/day | 0.323 mg/kg/day |
| hERG I inhibitor | No | No | No | No |
| hERG II inhibitor | No | No | No | No |
| Oral Rat Acute Toxicity (LD50) | 2.158 mol/kg | 2.285 mol/kg | 2.455 mol/kg | 2.097 mol/kg |
| Oral Rat Chronic Toxicity (LOAEL) | 1.133 mg/kg | 1.768 mg/kg | 2.409 mg/kg | 2.096 mg/kg |
| Hepatotoxicity | No | No | No | No |
| Skin Sensitisation | No | No | No | No |
| *T.Pyriformis* toxicity | 0.495 ug/L | 0.314 ug/L | 0.326 ug/L | 0.431 ug/L |
| Minnow toxicity | 1.951 mM | 2.36 mM | 3.169 mM | 0.483 mM |

### 3.5. Prediction of drugs target categories and structurally related analogs using DrugBank

Enzymes, kinase proteins, and oxidoreductases (such as aldose reductase and aldo-keto reductase) accounted for the majority of the target class (**Fig 7**). Compared to the other two compounds (Brevelin A and Pectolinarigenin), the target class Luteolin was predicted with a significantly greater percentage of probability (100%) (**Table 5**).

**Fig 7.**
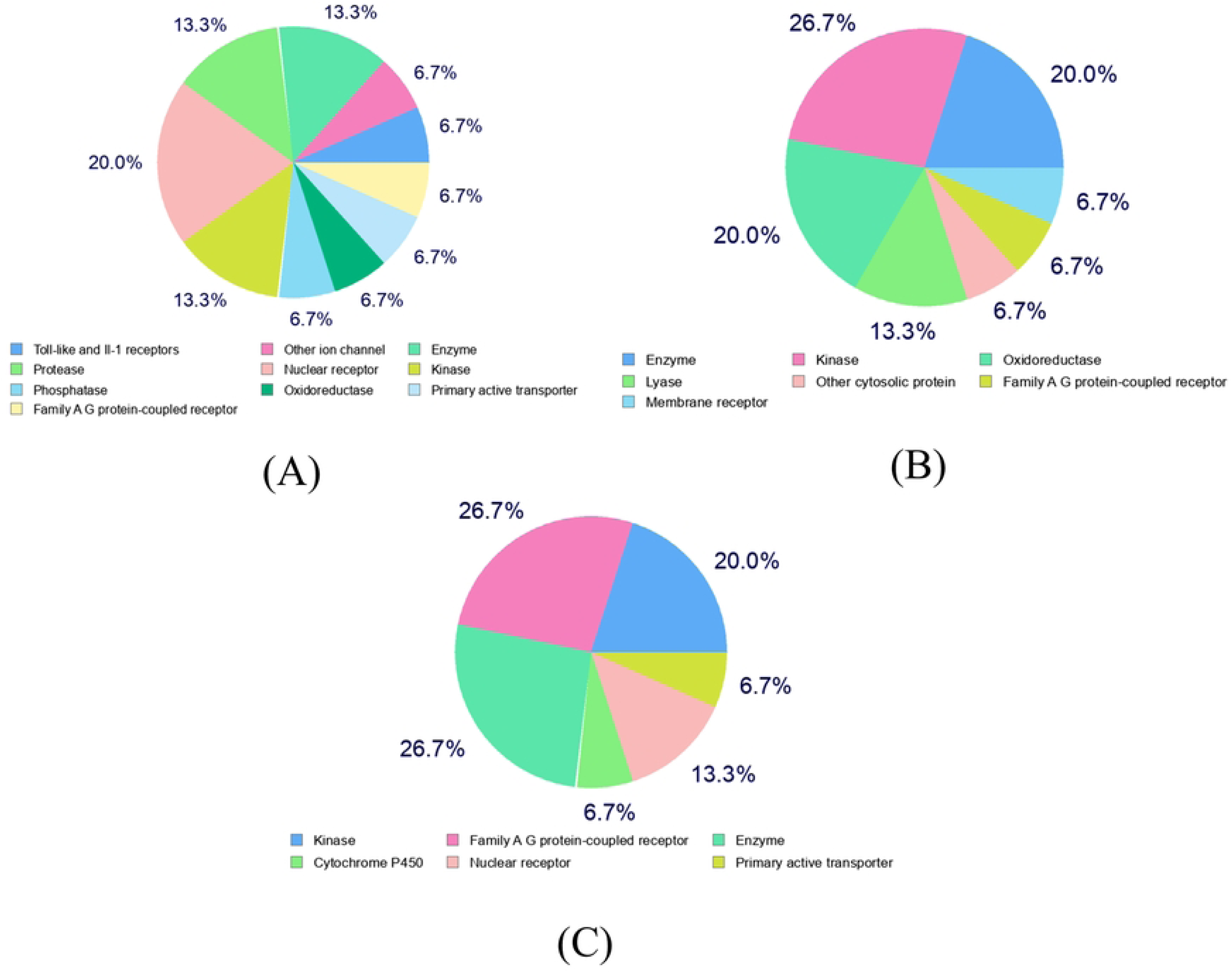
Prediction of target class for the top three lead compounds (A) Brevelin A, (B) Luteolin and (C) Pectolinarigenin.

**Table 5.**
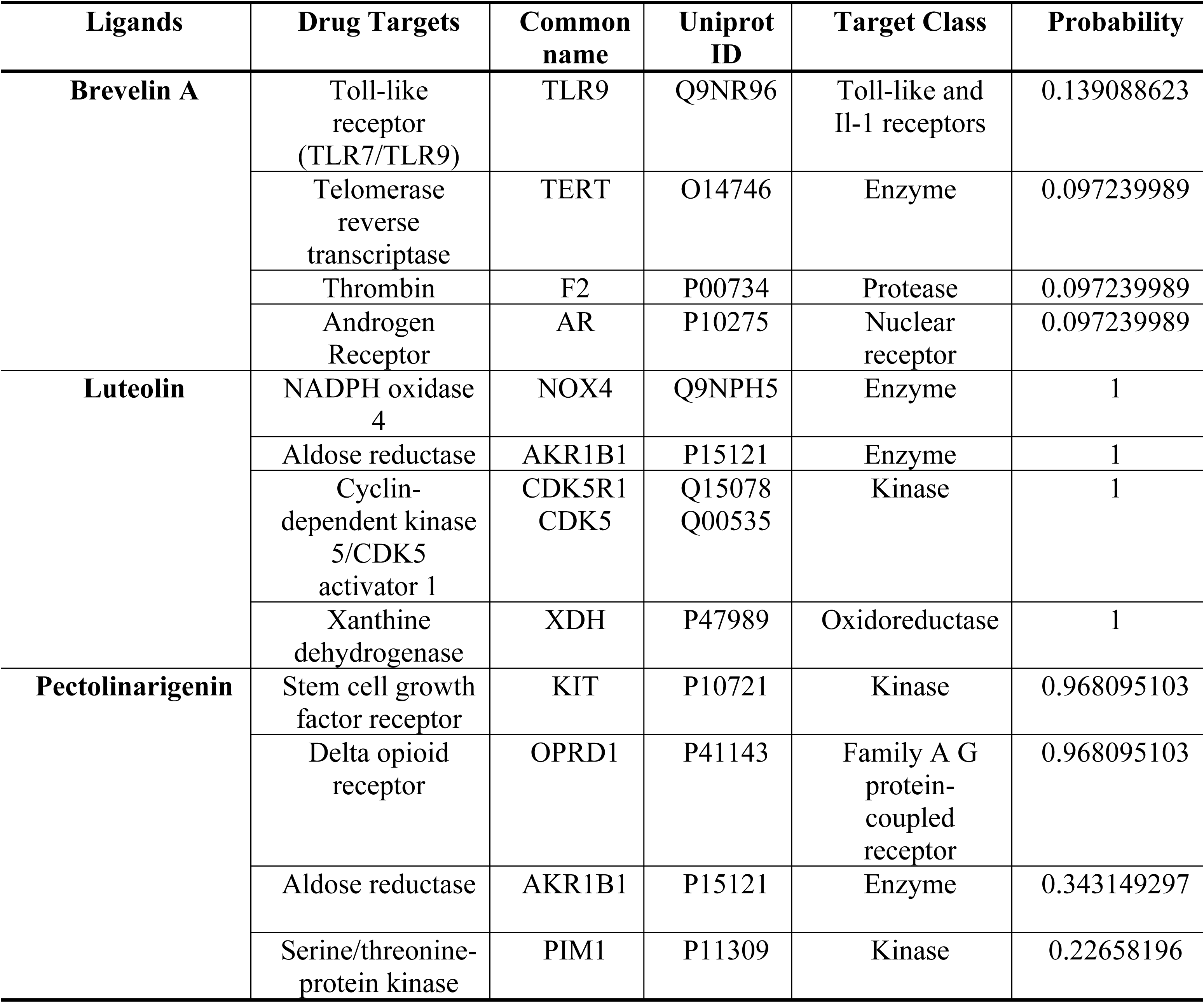
Predicted drug targets for Brevelin A, Luteolin and Pectolinarigenin. (By Swiss Target Prediction)

| Ligands | Drug Targets | Common name | Uniprot ID | Target Class | Probability |
| --- | --- | --- | --- | --- | --- |
| **Brevelin A** | Toll-like receptor (TLR7/TLR9) | TLR9 | Q9NR96 | Toll-like and Il-1 receptors | 0.139088623 |
|  | Telomerase reverse transcriptase | TERT | O14746 | Enzyme | 0.097239989 |
|  | Thrombin | F2 | P00734 | Protease | 0.097239989 |
|  | Androgen Receptor | AR | P10275 | Nuclear receptor | 0.097239989 |
| **Luteolin** | NADPH oxidase 4 | NOX4 | Q9NPH5 | Enzyme | 1 |
|  | Aldose reductase | AKR1B1 | P15121 | Enzyme | 1 |
|  | Cyclin-dependent kinase 5/CDK5 activator 1 | CDK5R1<br>CDK5 | Q15078<br>Q00535 | Kinase | 1 |
|  | Xanthine dehydrogenase | XDH | P47989 | Oxidoreductase | 1 |
| **Pectolinarigenin** | Stem cell growth factor receptor | KIT | P10721 | Kinase | 0.968095103 |
|  | Delta opioid receptor | OPRD1 | P41143 | Family A G protein-coupled receptor | 0.968095103 |
|  | Aldose reductase | AKR1B1 | P15121 | Enzyme | 0.343149297 |
|  | Serine/threonine-protein kinase | PIM1 | P11309 | Kinase | 0.22658196 |

To anticipate small compounds from DrugBank that are biologically active, ligand-based virtual screening was used. While Gibberellin A4 (DB07815) was found to be analogueous to Vernodalin, Gibberellic acid (DB07814) was found to be structurally similar to Brevelin A. Similarly, it was predicted that Hispidulin (DB14008) and Tricetin (DB08230) were analogous to Luteolin and Pectolinarigenin, respectively. Compared to Brevelin A and Vernodalin, the predicted analogs of luteolin and pectolinarigenin had the highest probability. The findings suggest that these could be potential drug candidates against cryptosporidiosis, thus requiring further experimental trials (**Table 6**). Brevelin A and Vernodalin scored lower on the probability scale than the predicted analogs of luteolin and pectolinarigenin. The results indicate that these could be promising therapeutic options for cryptosporidiosis, demanding additional testing.

**Table 6.**
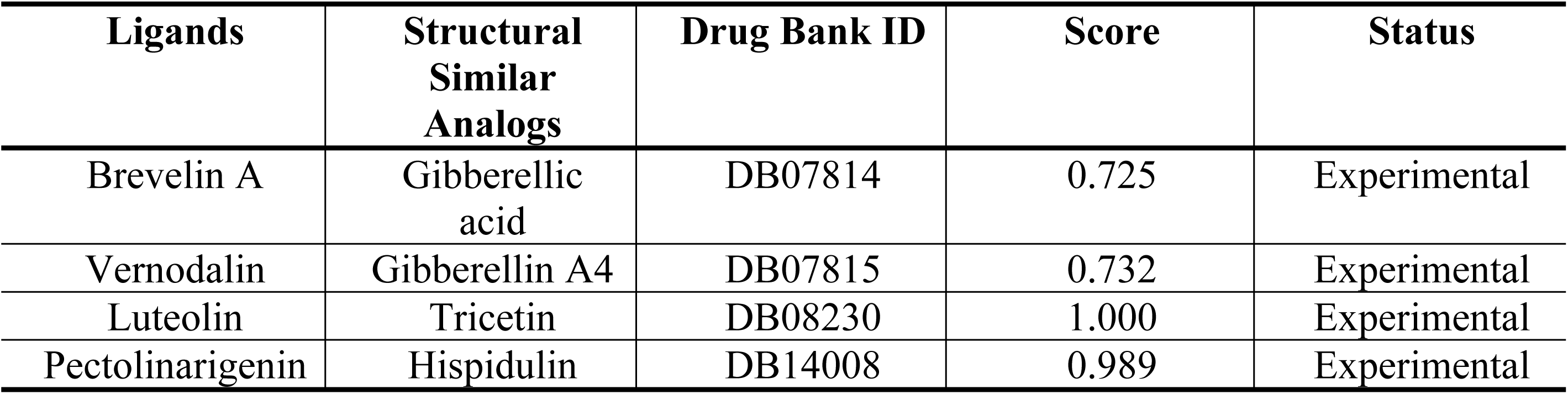
Predicted structurally similar compounds from Drugbank.

| Ligands | Structural Similar Analogs | Drug Bank ID | Score | Status |
| --- | --- | --- | --- | --- |
| Brevelin A | Gibberellic acid | DB07814 | 0.725 | Experimental |
| Vernodalin | Gibberellin A4 | DB07815 | 0.732 | Experimental |
| Luteolin | Tricetin | DB08230 | 1.000 | Experimental |
| Pectolinarigenin | Hispidulin | DB14008 | 0.989 | Experimental |

## 4. Discussion

The most frequent causes of the intestinal diarrheal illness cryptosporidiosis are *Cryptosporidium parvum* and *hominis.* These protozoan parasites are extensively present in both developed and developing countries, and they are a primary contributor to children’s malnourishment and severe diarrhea. In adults with healthy immune systems, infection is self-limiting; in immunocompromised people and children, however, it can become chronic. Drinking and recreational waters can become infected through the fecal oral pathway. Conventional techniques of treating water, such as filtration and bleaching, are ineffective against the infectious oocysts [51–53]. Because of this, researchers are always looking for suitable and effective drugs to treat cryptosporidiosis. As a result, utilizing a computational structure-based drug design strategy, this study tested 24 natural phytochemicals produced by various medicinal plants to determine their efficacy as inhibitory compounds against the Inosine Monophosphate Dehydrogenase protein from Cryptosporidium parvum.

The parasite *Cryptosporidium* is intracellular. These parasites can’t produce purine nucleotides de novo; instead, they must salvage adenosine from their host, according to a genomic study. A simplified process that turns adenosine into guanine nucleotides uses inosine 5′-monophosphate dehydrogenase (IMPDH) to catalyze the conversion from inosine 5′-monophosphate (IMP) to xanthosine 5′-monophosphate (XMP). In terms of structure, the parasite enzyme (CpIMPDH) differs from IMPDHs found in mammals. It indicates that lateral gene transfer occurred between bacteria and Cryptosporidium to acquire the IMPDH gene. As a result, CpIMPDH has become a desirable molecular target for developing effective drugs to treat diseases linked to this parasite [54–56]. Structure-based virtual screening, a computational biological method, has become an essential component of the biopharmaceuticals industry by speeding up the drug discovery process while saving money and time by visualizing the relationship of small molecules of interest or binding agents with macromolecules or receptors using the molecular docking method [57].

In this study, we have screened different phytochemicals that show potential activity against *Cryptosporidium*. Natural antioxidants like luteolin have been identified in a wide range of plants, including *Perilla frutescem (L.), Capsicum annuum L., and Ghrysanthemum indicum L*. It is anti-inflammatory, anti-oxidative, anti-tumor, and anti-viral, among many other advantageous properties. Lutein has been shown to suppress the inflammatory response by lowering the release of pro-inflammatory factors like interleukin 6 (IL-6) and tumor necrosis factor α (TNF-α) [58–60]. A study utilizing computational docking was conducted to examine luteolin’s inhibitory capacity. They discovered that luteolin inhibits the EGFR, or epidermal growth factor receptor. In order to combat lung cancer with chemotherapy, it is essential to find new, safe, and potent compounds due to the potentially fatal side effects of the recent EGFR mutant inhibitors, which include rash eruptions on the face, chest, back, and even the trunk, diarrhea, nausea, vomiting, anorexia, and stomatitis. Tumor-associated EGFR is present in around 10% of US patients with non-small cell lung cancer (NSCLC) and approximately 35% of patients in East Asia. These mutations affect EGFR exons 18–21, which encode a section of the kinase domain of EGFR and allow scientists to find drugs that exclusively bind and recognize cancer cells. As a result, EGFR mutations serve as both biomarkers and logical targets for focused treatment. Because of this, they use computational approaches to outsource the search for the best-in-class inhibitor for this druggable target [61].

Another molecular docking research demonstrated that luteolin shows antidiabetic effect by reversing hyperlipidemia, oxidative stress, and proinflammatory state in diabetic groups. As a result, luteolin may be an effective treatment or management option for diabetes [62]. Medicinal herbs have been the focus of research efforts in the search for new, effective antidiabetic drugs with minimal adverse effects. Because they have fewer or no adverse effects than synthetic medications, plant-derived phytoconstituents have drawn a lot of interest and are widely used in the primary healthcare systems of several developed and developing countries for the treatment of different diseases including diabetes [63].

In this study, Luteolin was found as a potential lead compound. It has a binding affinity of −8.6 kcal/mol with the 4IXH protein. Using molecular docking, ADMET (Absorption, Distribution, Metabolism, Excretion, and Toxicity) analyses, and molecular dynamics simulation, 25 bioactive phytochemicals of *C. asiatica* have been virtually screened to find potential anti-diarrheal agents. According to this study, one of the main compounds against the co-regulated pilus virulence regulatory protein (ToxT) of *V. cholerae* Toxin is Luteolin [64].

The development of innovative therapeutic drugs for managing a wide range of serious illnesses, such as cancer, diabetes, and inflammation, may find success by using of conventional medicinal plants in the coming years. There are several published reports on the biological evaluation and isolation of pectolinarigenin within the large family of flavonoids. These metabolites have demonstrated antimicrobial, antioxidant, anti-inflammatory, antidiabetic, and antitumor properties, making them a new target for a thorough comprehension of the mechanism of action [65]. Many different parasites use humans as their hosts in order to evolve, usually without causing immediate death to their host. Most parasites are undesirable or have the potential to impair human health, but certain parasitic illnesses (such as malaria and trypanosomiasis) can be lethal if not treated with suitable medications. Many flavonoid varieties have recently been found to be antiparasitic agents in plant extracts; however, an in-depth study of their structure-activity relationships (SARs) has yet to be completed [66, 67].

It has been studied how certain compounds from Baccharis uncinella’s aerial parts inhibit leishmaniasis. Pectolinarigenin demonstrated the strongest inhibition of the intracellular forms of both Leishmania (V.) braziliensis and Leishmania (L.) amazonensis, and it had significant effect on Leishmania (V.) braziliensis [68]. Flavonoids exhibit antiparasitic properties toward various organisms. They have been reported to bind to the nucleotide-binding site of MDR proteins, resulting in increased intracellular drug accumulation, making them a promising novel category of modulators for *Leishmania sp*. Furthermore, flavones enhance the effects of berbine, norfloxacin, and artemisinin on *Staphylococcus aureus* and *Plasmodium falciparum*, indicating a possible role for flavones in combination chemotherapy [69, 70]. In this present study, we have identified pectolinarigenin as a potential drug candidate against *Cryptosporidium parvum*. It has seven hydrogen bonds and three other hydrophobic interactions with the 4IXH protein.

Several sesquiterpene lactones isolated from different African Vernonia species have shown significant antischistosomal, plasmodicidal, and leishmanicidal action in vitro. The leaves of this species are often used in decoction form for diverse purposes in traditional medicine, including fevers, coughing, diarrhea, boils, and general tonicity [71]. The antibacterial compound vernodalin was extracted from the leaves of *V. colorata*, which demonstrated strong antibacterial activity in the disc-diffusion bioassay, according to a study [72]. Vernodalin has been found to have antitrypanosomal activity in another investigation [73]. In our present study, we revealed that vernodalin is a potent drug of choice against cryptosporidiosis. It has a binding affinity of −8.7 with the 4IXH protein, and it has seven ligand binding sites. Vernodalin also triggers the death of gastric cancer (GC) cells and suppresses the growth, adhesion, and metastasis of GC cells. It reduces tumor development and enhances apoptosis of gastric cancer cells by inhibiting the FAK/PI3K/AKT/mTOR and MAPK signaling pathways [74].

Sesquiterpene lactone Brevilin A was extracted from *Centipeda minima* and has the ability to suppress the proliferation of many types of tumor cells. In vivo and in vitro colon adenocarcinoma CT26 cell proliferation was examined in this work, as well as the molecular mechanism underlying Brevilin A’s inhibitory impact. The outcomes showed that apoptosis was the cause of Brevilin A’s dose-dependent inhibitory effect on CT26 proliferation. Moreover, Brevilin A dose-dependently raised ROS levels, lowered mitochondrial membrane potential (MMP), and triggered CT26 cell death. According to a study, Brevilin A’s anti-tumor efficacy was mostly attributed to the stimulation of cell apoptosis and autophagy, indicating a bright future for the medication as an anticancer treatment for colon adenocarcinoma [75].

Another investigation was done on aberrant signals in human cells, which typically have unexpected adverse impacts on the health of an individual. They concentrate on these forms of JAK-STAT signal pathway events, particularly those that are brought on by aberrantly activated STAT3, an oncoprotein that plays a crucial role in immunological disorders and many tumor types’ processes of cell survival, growth, and proliferation. Through the development of a high-throughput drug screening system based on the STAT3 signal in human lung cancer A549 cells, they have evaluated a natural product library containing compounds that have been isolated from medicinal herbs. Brevilin A, one of the compounds, demonstrated both substantial inhibition of the STAT3 signal and inhibition of cell growth that was dependent on the STAT3 signal. Subsequent research showed that Brevilin A suppresses both STAT1 and STAT3 signaling, which phosphorylates STAT3 and STAT1 and increases the expression of their target genes when cytokines are present. Furthermore, they discovered that Brevilin A might inhibit JAK’s tyrosine kinase domain, JH1, which would reduce JAK activity. Brevilin A dramatically inhibited cytokine-induced phosphorylation of STATs and other substrates. The way that Brevilin A targets JAK activity suggests that it can be employed as a drug that blocks other JAK-STAT hyperactivation in addition to its potential utility as a STAT3 inhibitor. Therefore, these results strongly encouraged the development of therapeutic medicines and specific JAK-STAT inhibitors to increase the survival rate of patients with hyperactivated JAKs and STATs [76]. A molecular docking study also revealed that Brevilin A has anticancer activity [77].

The potentially fatal illness known as acute lung injury (ALI) is typified by an uncontrollably high inflammatory response. As the primary kinases in the NF-κB pathway’s activation, IKKα/β are linked to inflammatory lung damage and make appealing targets for ALI treatment. The Chinese herb Centipeda minima yields a sesquiterpene lactone called Brevilin A (BVA), which is used to treat inflammatory conditions [78]. In this study, it was discovered that Brevilin A has a binding affinity of −8.9 with *Cryptosporidium parvam’*s protein (4IXH), which indicates that it could be a potential drug of choice in the case of cryptosporidiosis treatment. We found that it has three ligand binding sites (Thr22, Gly214, and Lys210) and four hydrogen bonds with the 4IXH protein. A research idea was designed to assess if and to what extent an inhibitor’s apparent anti-cryptosporidial efficacy was due to its impact on the parasite target. The host cells with transient overexpression of the multidrug resistance protein-1 (MDR1) showed significantly increased drug tolerance. The transient transfection paradigm, however, could only be employed to evaluate native MDR1 substrates. This group also reported on another study in which they used a stable MDR1-transgenic HCT-8 cell model, an advanced model that enables multiple rounds of drug selection to rapidly create novel resistance to non-MDR1 substrates. The sole FDA-approved drug for treating human cryptosporidiosis, nitazoxanide, was able to totally kill *C. parvum* by addressing the parasite target, as demonstrated by their successful validation of this finding using the new model [79]. In our present study, we used nitazoxanide as a control. By comparing it with this control drug, Brevelin A, Vernodalin, Luteolin, and Pectolinarigenin were revealed as new potential drug candidates against the *C. parvum* parasite.

We used *in silico* screening to screen a total of 24 naturally occurring bioactive phytochemicals in this study. Excellent biochemical interactions involving hydrogen bonds and hydrophobic interactions were revealed by the molecular docking result between the ligands and the receptor. Additionally, in comparison to the reference drugs, nitazoxanide, which has a binding affinity of - 8.1 kcal/mol, some of the inhibitors showed superior affinities that ranged from −8.1 to −8.9 kcal/mol. In silico pharmacokinetics properties and ADMET predictions were examined in order to assess the drug-like profiles and pharmacological characteristics of the targeted inhibitors. Lipinski’s rule of five is not broken by any of the targeted ligands, including the control drug, according to the computed physicochemical parameters. This indicates that all of the inhibitors have the potential to be oral bioavailable and pharmacologically active drugs. Similarly, the ADMET characteristics showed that all of the inhibitors have good characteristics for excretion, metabolism, absorption, and distribution; none of the inhibitors are toxic or carcinogenic to humans. In a similar vein, the BOILED-Egg graphics show that every ligand has a high probability of being absorbed by the human gastrointestinal system and possibly penetrating the brain.

## 5. Conclusion

No entirely effective therapies are available for treating the moderate-to-severe, occasionally fatal, watery diarrhea caused by the zoonotic protozoan parasite *Cryptosporidium parvum.* The notion of utilizing natural products for drug discovery has been reinvigorated in order to mitigate the deleterious effects of synthetic drugs. Natural plant-based products offer a wealth of benefits in the fight against numerous pathogenic illnesses and diseases, even with their few disadvantages. According to this study, all of the chosen inhibitors—Brevelin A, Vernodalin, Luteolin, and Pectolinarigenin—would be highly active as potential drugs based on predicted bioactive parameters. It is highly recommended to conduct extensive *in vitro* studies to confirm our *in silico* outcomes.

## Author Contributions

**Conceptualization:** Abdullah Al-Mamun, Mohabbat Hossain, Nazneen Naher Islam

**Formal analysis**: Abdullah Al-Mamun, Shah Imran Hossain

**Methodology:** Abdullah Al-Mamun, Shah Imran Hossain, Abu Tayab Moin, Md Shafiqul Islam Rakib, Md Mahedi Hasan, Eshma Binte Yousuf, Shams Nur Powshi, Enayetul Islam

**Project administration**: Mohabbat Hossain, Nazneen Naher Islam

**Supervision**: Dil Umme Salma Chowdhury, Mohabbat Hossain, Nazneen Naher Islam

**Visualization**: Abdullah Al-Mamun, Shah Imran Hossain, Abu Tayab Moin

**Writing – original draft:** Abdullah Al-Mamun, Shah Imran Hossain, Mohabbat Hossain

**Writing – review & editing:** Nasrin Jahan Salma Tumpa, Asmaul Hosna, Dil Umme Salma Chowdhury, Mohabbat Hossain, Sabrina Shameen Alam, Nazneen Naher Islam

